# Rapid evolution of primate type 2 immune response factors linked to asthma susceptibility

**DOI:** 10.1101/080424

**Authors:** Matthew F. Barber, Elliot M. Lee, Hayden Griffin, Nels C. Elde

## Abstract

Host immunity pathways evolve rapidly in response to antagonism by pathogens. Microbial infections can also trigger excessive inflammation that contributes to diverse autoimmune disorders including asthma, lupus, diabetes, and arthritis. Definitive links between immune system evolution and human autoimmune disease remain unclear. Here we provide evidence that several components of the type 2 immune response pathway have been subject to recurrent positive selection in the primate lineage. Notably, rapid evolution of the central immune regulator *IL13* corresponds to a polymorphism linked to asthma susceptibility in humans. We also find evidence of accelerated amino acid substitutions as well as repeated gene gain and loss events among eosinophil granule proteins, which act as toxic antimicrobial effectors that promote asthma pathology by damaging airway tissues. These results support the hypothesis that evolutionary conflicts with pathogens promote tradeoffs for increasingly robust immune responses during animal evolution. Our findings are also consistent with the view that natural selection has contributed to the spread of autoimmune disease alleles in humans.

## INTRODUCTION

Executing a balanced immune response is essential for organism survival. While robust immune activation is required to adequately recognize and clear infections, excessive inflammation contributes to a wide variety of disease conditions including arthritis, diabetes, asthma, and septic shock. Asthma is of particular concern as the incidence among humans has increased dramatically over the past half century (Akinbami et al. 2016). While exhibiting markedly variable disease pathology, asthma is generally characterized by chronic inflammation of the conducting airways leading to hypersensitivity (AHR) (Lambrecht and Hammad 2012). Work over the last several years greatly advanced our understanding of the genetic and physiologic processes contributing to asthma. It is now appreciated that the type 2 immune response is a major contributor to this disease in many patients (Figure 1A) (Holgate 2012). Recognition of allergens by antigen presenting cells leads to maturation of T helper 2 (Th2) cells and production of associated cytokines as well as proliferation of eosinophils, basophils and mast cells that contribute to AHR.

Eosinophils are a class of granulocytes that undergo activation and expansion in response to Th2-mediated immune signaling (Hogan et al. 2008). These cells are strongly recruited to sites of allergic inflammation, including the asthmatic airway. Eosinophil granules contain several highly cationic proteins with potent antimicrobial and anti-helminth activity, which are released at sites of infection (Acharya and Ackerman 2014). Key among these is the eosinophil major basic protein (MBP), which binds to microbial and host cell surfaces promoting membrane disruption and cell death. Eosinophil granules also contain high levels of eosinophil derived neurotoxin (EDN, also called RNASE2) and eosinophil cationic protein (ECP, also called RNASE3), two members of the RNaseA protein family. Eosinophil peroxidase (EPX) is a fourth major granule protein which catalyzes the production of reaction oxygen species to impair microbial growth. This combination of potent effectors make eosinophils key responders in the type 2 immune response. However, eosinophil granule proteins are also toxic to host cells and tissues, with experimental evidence indicating that these proteins contribute to asthma severity (Nielsen et al. 2009).

Despite progress in understanding the molecular mechanisms that underlie disease pathology, asthma remains a complex syndrome with a diverse spectrum of symptoms and severities. Previous genome-wide association studies (GWAS) implicate multiple underlying genetic risk factors, further highlighting the biological complexity of this disease (Tamari et al. 2013). While human population-based approaches have been repeatedly applied to understand asthma susceptibility, less attention has been given to species-level comparative genetic approaches.

Molecular phylogenetics provides a useful framework from which to analyze millions of years of genetic variation to investigate mechanisms underlying diverse molecular and biological phenomena (Dean and Thornton 2007; Harms and Thornton 2013). Immune system components are some of the most rapidly evolving genes in vertebrates, as host populations continually adapt against infectious pathogens (George et al. 2011; Rausell and Telenti 2014). Previous studies have used phylogenetic signals of rapid evolution to dissect molecular features of evolutionary “arms races” unfolding at host-microbe interfaces (Sawyer et al. 2005; Elde et al. 2009; Barber and Elde 2014). Such evolutionary approaches can further guide functional studies probing the basis for microbial host range and mechanisms of protein evolution (Daugherty and Malik 2012). Host genes involved in recurrent genetic conflicts are often characterized by strong signatures of positive selection, reflecting transient advantages of novel genetic variants during evolution with microbes. Much of the work in this field has focused on implications for rapid evolution on host defense and infectious disease susceptibility (Lim et al. 2012; Mitchell et al. 2012).

Adaptation in response to pathogens may also incur a cost to the host, as documented in the case of sickle cell disease where particular alleles provide enhanced resistance to malaria at the cost of severe anemia in homozygous carriers (Kwiatkowski 2000). Recent work has also suggested that rapid evolution of immune system components can contribute to the establishment of species barriers by producing lethal genetic incompatibilities (Chae et al. 2014). In the present study we identify signatures of positive selection in primate immune genes that correspond to asthma susceptibility loci in human populations, suggesting that host-microbe evolutionary conflicts contribute to the spread or persistence of autoimmune disease susceptibility alleles.

## MATERIALS & METHODS

### Primate genetic sources

Genomic DNA and total RNA were obtained from primary cells (Coriell Cell Repositories) from the following species: *Homo sapiens* (human; primary human foreskin fibroblasts; gift from A. Geballe), *Pan troglodytes* (chimpanzee; PR00226), *Gorilla gorilla* (western lowland gorilla; AG05251), *Pongo pygmaeus pygmaeus* (Bornean orangutan; AG05252), *Hylobates lar* (white-handed gibbon; PR01131), *Hylobates leucogenys* (white-cheeked gibbon; PR00712), *Hylobates syndactylus* (island siamang; PR00722), *Macaca mulatta* (rhesus monkey liver tissue; gift from S. Wong), *Papio anubis* (olive baboon; PR00036), *Lophocebus albigena* (black crested mangabey; PR01215), *Allenopithecus nigroviridis* (Allen’s swamp monkey; PR01231), *Cercopithecus aethiops* (African green monkey; PR01193), *Erythrocebus patas* (patas monkey; NG06254), *Cercopithecus mitis* (blue monkey; PR00996), *Cercopithecus Wolfi* (Wolf’s guenon; PR00486), *Colobus guereza* (colobus monkey; PR00240), *Saguinus fuscicollis* (Spix’s saddle-back tamarin; AG05313), *Sanguinus labiatus* (mustached tamarin; AG05308), *Callithrix geoffroyi* (white-fronted marmoset; PR00789), *Lagothrix lagotricha* (common woolly monkey; AG05356), *Saimiri sciureus* (common squirrel monkey; AG05311), *Aotus nancyma* (night monkey; PR00627), *Callicebus moloch* (dusky titi; PR00742), *Alouatta sara* (Bolivian red howler monkey; PR00708), *Pithecia pithecia* (white-faced saki; PR00239). Additional sequences were obtained from NCBI GenBank entries.

### Gene sequencing and cloning

Genomic DNA and total RNA were harvested using the AllPrep DNA/RNA Mini kit (Qiagen). Isolated RNA (50 ng) from cell lines was used as template for RT-PCR (SuperScript III; Invitrogen). PCR products were TA-cloned into pCR2.1 (Invitrogen) and directly sequenced from at least three individual clones.

### Phylogenetic and protein structure analyses

DNA multiple sequence alignments were performed using MUSCLE, and sequences were trimmed manually. Unless otherwise noted, primate species phylogenies were used for downstream evolutionary analysis (Perelman et al. 2011). Alternatively, maximum-likelihood gene phylogenies were generated using PhyML with SPR topology search and bootstrapping for branch support (Guindon et al. 2010). Substitution models were chosen based on ProtTest algorithm (Abascal et al. 2005). Tests for positive selection were performed using codeml from the PAML software package with both F61 and F3X4 codon frequency models. A free-ratio model (M0) allowing *d*N/*d*S (**ω**) to vary along branches of the phylogeny was used to calculate *d*N/*d*S values between lineages (Yang 2007). Positive selection was assessed by fitting the multiple alignment to either F3x4 or F61 codon frequency models. Likelihood ratio tests (LRTs) were performed by comparing pairs of sitespecific models (NS sites): M1 (neutral) with M2 (selection), M7 (neutral, beta distribution of *d*N/*d*S<1) with M8 (selection, beta distribution, *d*N/*d*S>1 allowed). Alternative tests which also account for synonymous rate variation and recombination, including FUBAR, FEL, and MEME, were performed using the HyPhy software package (Pond et al. 2005; Delport et al. 2010). The BUSTED algorithm was employed to test for gene-wide signatures of episodic positive selection. Sites under positive selection were mapped onto three-dimensional molecular structures using Chimera (Pettersen et al. 2004)(http://www.cgl.ucsf.edu/chimera/).

## RESULTS

### Signatures of positive selection among primate type 2 immune response genes

To assess whether allergic immunity factors have been subject to positive selection in primates, we first compiled orthologs of 17 genes which represent a subset of the core type 2 immune response (Figure 1A, Supplementary Figure 1). This gene set includes numerous cytokines that mediate Th2 cell signaling associated with asthma (*IL3*, *IL4*, *IL5*, *IL9*, *IL10* and *IL13*), as well as components of IgE signaling (*FCERA-C*) and eosinophil granule proteins (*ECP*, *EDN*, *EPX,* and *MBP*). For each gene, nine orthologs from anthropoid primates were obtained from available public databases. *ECP* (also known as *RNASE3*) was the one exception, as this gene arose via a recent duplication of *EDN* (also known as *RNASE2*) in the common ancestor of Old World monkeys and apes, and is thus absent from New World monkey genomes. Orthologous protein-coding regions were aligned, manually trimmed, and phylogenetic analysis was performed to screen for evidence of positive selection using the PAML software package (NS sites), which infers signatures of selection from ratios of non-synonymous to synonymous substitution rates (termed *d*N/*d*S, or **ω**). Of the 17 genes selected, six (*ECP, MBP, IL13, IL3, FCER1A*, and *EDN*) exhibited signatures of positive selection (*P*<0.01 based on M7 vs. M8 likelihood ratio test statistics) (Figure 1B). These results indicate that several components of the type 2 immune response pathway may have been subject to positive selection in anthropoid primates.

**Figure 1.**
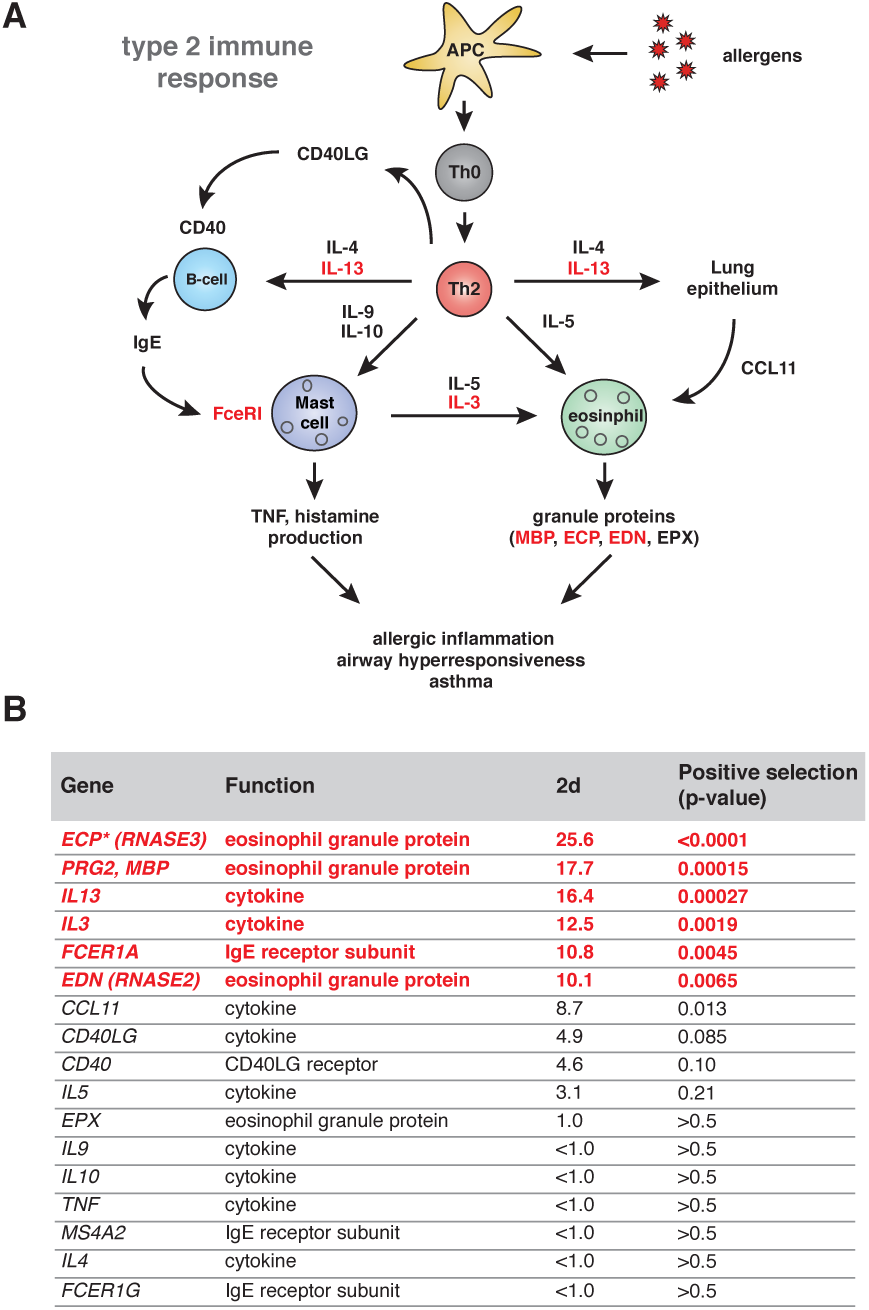
Signatures of positive selection in the primate type 2 immune response pathway. **A.** Overview of the type 2 immune response pathway in humans. Recognition of allergens by antigen presenting cells (APCs) leads to activation of naïve Th0 cells into mature Th2 cells. Secreted cytokines IL-13, IL-4, IL-5, IL-9 and IL-10 contribute to activation of B-cells, granulocytes and remodeling of the lung epithelia. Together these responses contribute to allergic inflammation characteristic of asthma in conducting airways. Genes exhibiting signatures of positive selection are shown in red. **B.** Summary of screen for evidence of positive selection among allergic immune response factors. Nine primate orthologs for each gene representing hominoid, Old World, and New World monkeys were analyzed using codeml from the PAML software package. P-values and 2d statistics are shown for model comparisons of M7 versus M8. Genes with statistical support (*P*<0.01) for positive selection are highlighted in red. \**ECP* is only present in hominoid and Old World monkey genomes, preventing the inclusion of New World monkeys for phylogenetic analyses.

### Variation in interleukin 13 among primates corresponds to a human asthma susceptibility allele

The group of type 2 immune response factors displaying support for positive selection included interleukin 13 (IL-13), a major regulator of the Th2 immune pathway. IL-13 is an inflammatory cytokine that promotes numerous aspects of allergic inflammation, including B cell class switching to stimulate IgE production, mast cell and eosinophil maturation, mucus production, and angiogenesis (Wills-Karp et al. 1998; Ingram and Kraft 2012). To improve our resolution of IL-13 evolution in primates, we cloned and sequenced additional *IL13* orthologs using genomic DNA from a diverse set of primate primary cell lines (Figure 2A, Supplementary Tables 1-3, 7-9). Phylogenetic analysis of the expanded dataset using PAML supported evidence of positive selection acting on *IL13* (*P* < 0.006), while BUSTED did not (*P* = 0.11). One explanation for this discrepancy could be a gene-wide low level of sequence variability observed between primate *IL13* orthologs, which might limit the statistical power of tests for selection based on relative substitution rates (Supplementary Figure 2, Supplementary Data). We therefore considered specific sites in IL-13 that contributed to signatures of positive selection.

**Figure 2.**
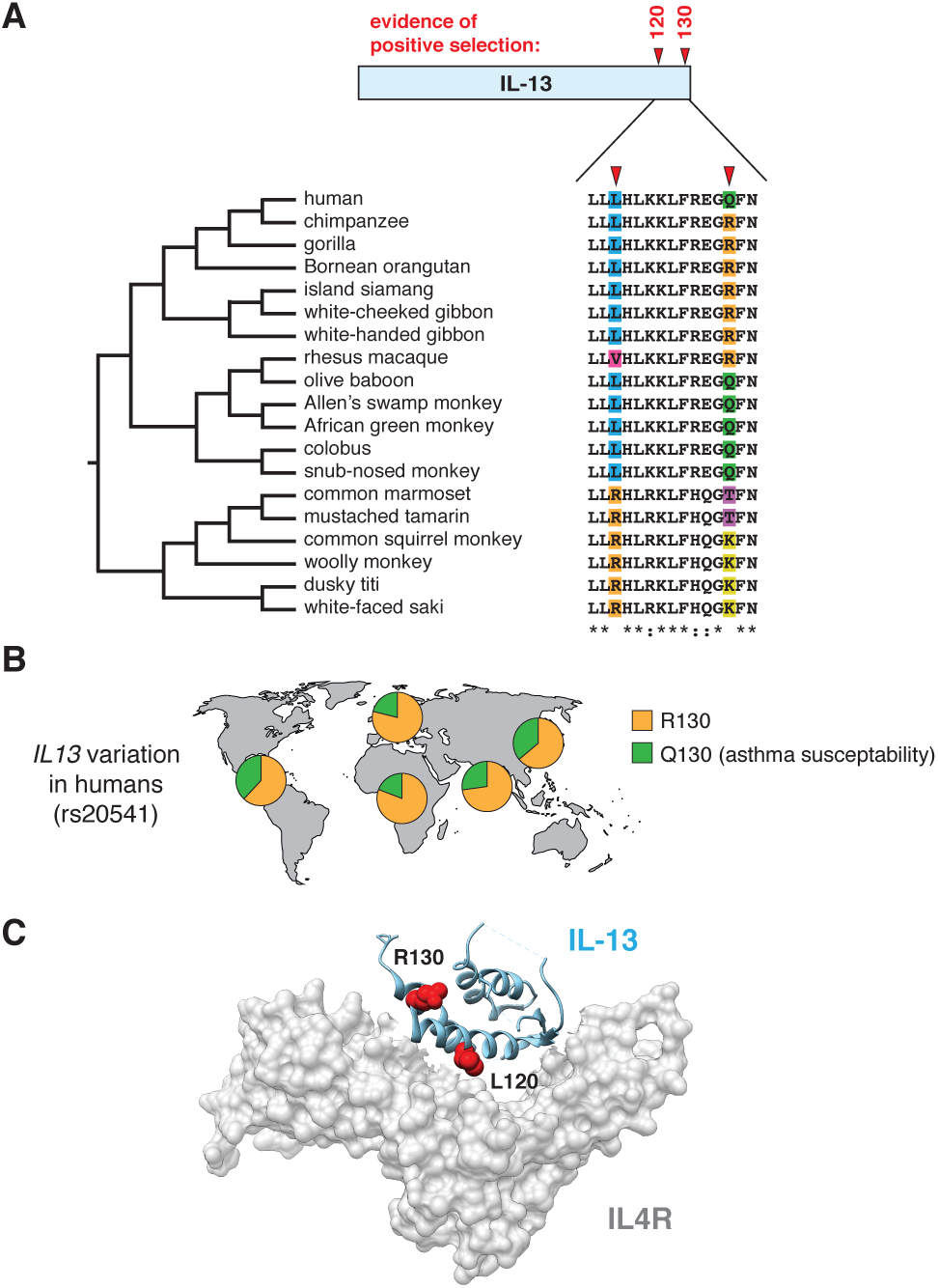
Variation in interleukin 13 among primates associated with human asthma susceptibility. **A.** Analyses using PAML and the FUBAR algorithms narrowed signatures of selection in primate IL-13 to two amino acid positions, 120 and 130. Amino acid variability at these sites across 19 primate species is highlighted. **B.** Schematic depicting allele frequencies for *IL13* R130 and Q130 variants across human populations. Individuals carrying the Q130 allele have previously been shown to have increased asthma susceptibility. Data from the 1000 Genomes Project (phase 3). **C.** Co-crystal structure (PDB: 3BPO) of IL-13 (blue) in complex with its receptor, IL4R (gray). Sidechains of amino acids subject to positive selection across primates are highlighted in red.

Using two independent algorithms (PAML and FUBAR), we identified two sites in the IL-13 C-terminus, amino acid positions 120 and 130, that display signatures of positive selection (Figure 2A). Notably, an abundant human polymorphism in *IL13* at position 130 has previously been linked to asthma susceptibility (Figure 2B) (Heinzmann 2000; Heinzmann et al. 2000; Wang et al. 2003). All hominoids surveyed contain arginine at position 130, while most Old World monkeys contain a glutamine at this site, corresponding to the ancestral and derived human alleles, respectively. Previous functional studies of human IL-13 indicate that the Q130 variant (present at roughly 25-30% allele frequency in humans) possesses enhanced downstream signaling functions, including increased IgE production (Vladich et al. 2005). New World monkeys contain both lysine and tyrosine residues at this position, not found in either hominoids or Old World monkeys. The impact of these alternative substitutions on downstream IL-13 signaling is presently unknown. We also observed that position 120 exhibits limited signatures of positive selection in primates, a site that lies directly in the binding interface of the IL-13 receptor (Figure 2C). As the relative affinity of IL-13 for its cognate receptor is critical for downstream signaling processes, we hypothesize that variation at this site could strongly impact IL-13 function. These results establish a genetic link between variation in primate *IL13* and human asthma susceptibility.

### Eosinophil major basic protein (MBP) has been subject to episodic positive selection

Three of the six rapidly evolving type 2 immune response genes identified in our initial screen form components of eosinophil granules (Figure 1B). MBP is one of the most highly cationic proteins in the human proteome, composed almost entirely of an atypical lectin-like domain. Like other lectin related proteins, MBP is capable of binding cell surface glycans such as heparin (Acharya and Ackerman 2014). Binding of MBP to cell surfaces, combined with its strong cationic charge and propensity to form oligomeric complexes, has been hypothesized to promote membrane disruption and toxicity against microbes as well as host cells. Based on evidence of positive selection from our initial screen, we expanded our analysis of MBP evolution in primates by cloning and sequencing additional orthologs using complementary DNA (cDNA) derived from primary primate cell lines. *MBP* possesses strong signatures of positive selection along several branches of the hominoid, Old World and particularly New World monkey phylogeny (Figure 3A, Supplementary Tables 1, 2, 4, 7-9). In contrast to IL-13, we observe that a high proportion of sites at diverse surfaces of MBP contribute to signals of positive selection. Eight of the nine most rapidly evolving sites lie in the C-terminal protein domain, indicating that selection has largely acted upon functions of the cleaved, mature form that is released from granules (Figure 3B, C). Recent work has also suggested that MBP forms an amyloid-like oligomeric structure on cell surfaces which facilitates membrane disruption (Soragni et al. 2015). Structural and computational predictions mapped this amyloid-forming domain to a small stretch of five amino acids on the exterior of the protein. One site within this “amyloid zipper,” position 134 (valine in humans) is among the nine sites with signals of positive selection in our sampling of primates (Figure 3B), suggesting that variation at this site could impact protein oligomer formation and target cell membrane disruption. Together these results suggest that rapid evolution of primate MBP modulates toxicity with functional consequences for host immunity.

**Figure 3.**
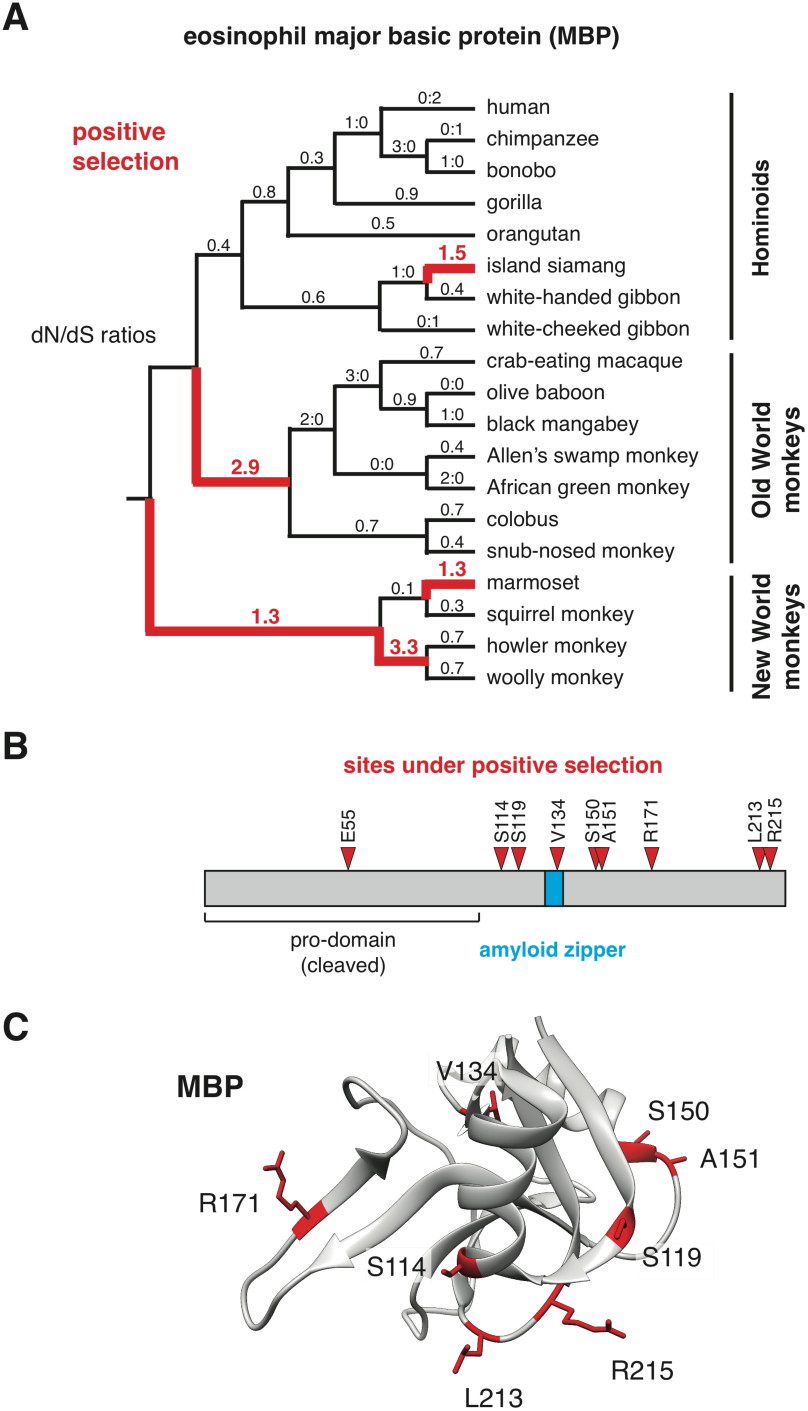
Rapid evolution of the eosinophil major basic protein. **A.** *MBP* gene was sequenced from the indicated primate species and *d*N/*d*S ratios were calculated across the primate species phylogeny using PAML. Lineages with elevated *d*N/*d*S (>1), suggesting of positive selection, are highlighted in red. In cases lacking either non-synonymous or synonymous mutations, ratios of respective substitution numbers are indicated. **B.** Diagram showing sites identified under positive selection in MBP (Pr>0.99, Naïve Empirical Bayes analysis in PAML). The positions of the prodomain, which is cleaved during protein processing, and the amyloid zipper which mediates protein oligomer formation, are indicated. **C.** Sites under positive selection identified in panel B, mapped onto the crystal structure of the mature MBP (PDB: 1H8U). Identified sites appear to occupy several distinct protein surfaces.

### Recurrent gain and loss of primate-specific eosinophil RNaseA genes

In addition to *MBP*, our evolutionary screen identified strong signatures of positive selection in *ECP*, another component of eosinophil granules. *ECP* is a comparatively young gene, emerging roughly 30 million years ago from a duplication of *EDN* in the common ancestor of hominoids and Old World monkeys (Zhang and Rosenberg 2002). ECP and EDN are members of the RNaseA protein family; however, experimental evidence suggests that RNase enzymatic activity has been subsequently lost or impaired in ECP (Zhang and Rosenberg 2002). Like other eosinophil granule proteins, ECP possesses a strong net cationic charge, which has been proposed to contribute to toxicity against target cells. ECP is also capable of interacting with surface glycans which can mediate its association with cell membranes, similar to MBP (Bystrom et al. 2011). A previous study concluded that *ECP* has been subject to strong positive selection since its divergence from *EDN* (Zhang et al. 1998), although these studies did not pinpoint specific sites in *ECP* that contribute to this signature.

Efforts to expand our dataset of *ECP* orthologs led to an unexpected finding – while several ECP transcripts were detected using PCR, we also amplified transcripts that appear distantly related from *bona fide ECP* orthologs. In particular, *ECP* genes cloned from bonobo and orangutan cDNA are markedly divergent from other hominoid *ECP* orthologs (Figure 4A, Supplementary Figure 3). Inferences of the primate *ECP*/*EDN* gene phylogeny revealed that these transcripts appear most similar to extant primate *ECP* genes, despite their sequence divergence. While surrounding DNA regions further suggest that these genes are true orthologs of *ECP*, we cannot fully exclude the possibility that these genes arose from distinct duplication events followed by gene loss of ancestral *ECP*. Notably, humans possess an *ECP*-related pseudogene, *ECRP*, proximal to both *ECP* and *EDN* on chromosome 14. The existence of such pseudogenes suggests that duplication events in this gene cluster could be ongoing in other primate taxa. In addition, gene phylogenies of several *ECP* orthologs cloned from Old World monkeys appear to break predictions based on species relationships (Figure 4A). Such discordance could be the result of intense episodes of positive selection, recent *ECP* gene duplication events, or incomplete lineage sorting.

In addition to evidence of gene duplication, we also observed instances of potential *ECP* gene loss in the *Hylobatidae* (gibbon) lineage. Multiple cloning efforts failed to yield *ECP* orthologs from three separate gibbon species using either cDNA or genomic DNA. We likewise failed to identify a clear *ECP* ortholog from the recently assembled white-cheeked gibbon genome using BLAST. Instead, the region of gibbon chromosome 22 which contains the ancestral *EDN* gene appears to have undergone a large-scale genomic rearrangement, with the segment containing *ECP* in other apes removed (Figure 4B). Such dynamic genome rearrangements have previously been documented in gibbon genomes (Carbone et al. 2014). We were also unable to isolate the *ECP*-containing region on other scaffolds of the gibbon genome assembly. These results suggest that *ECP* may have been lost or pseudogenized specifically in the *Hylobatidae* lineage during a chromosomal translocation event.

**Figure 4.**
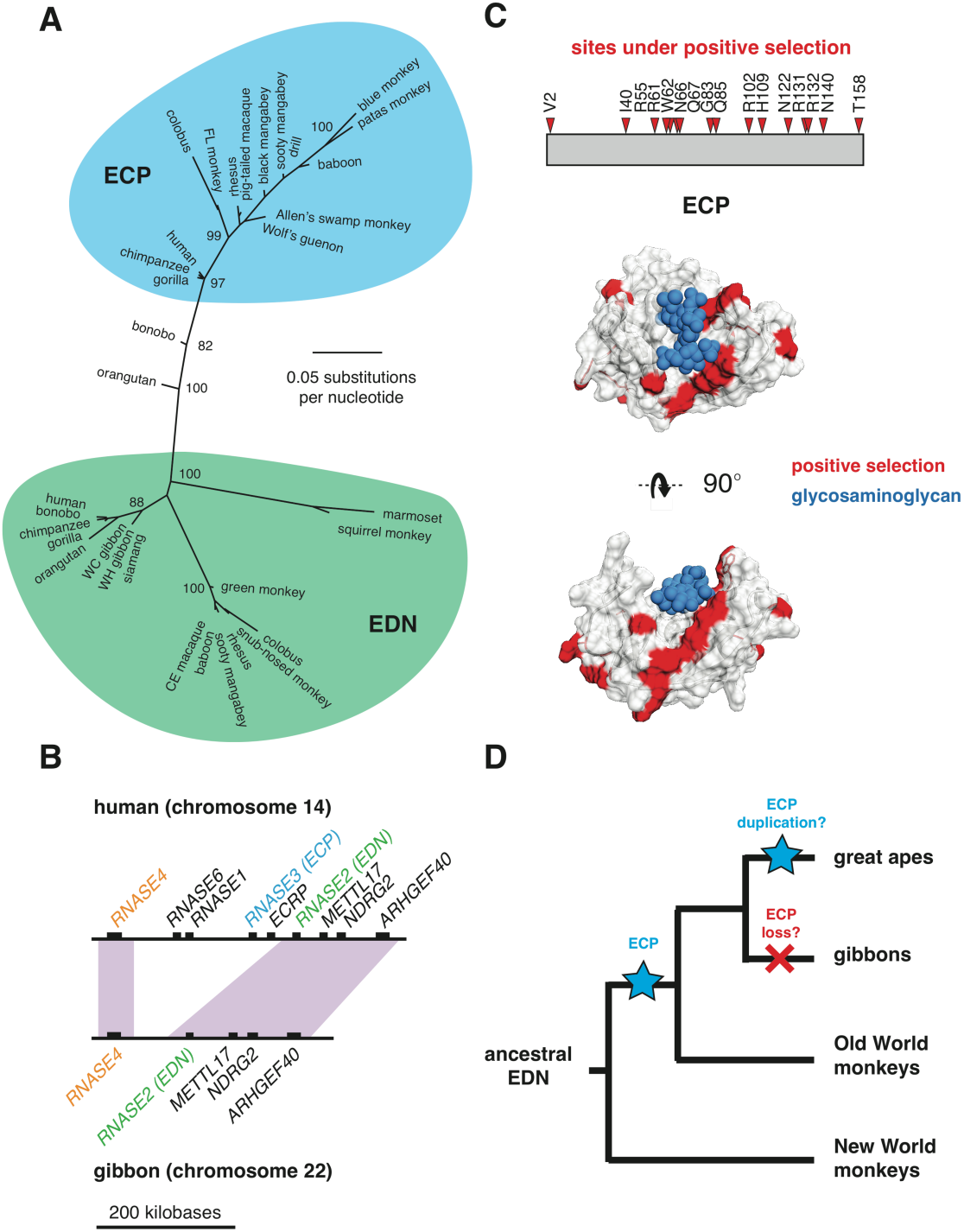
Recurrent gene gain, loss, and amino acid changes shape primate eosinophil RNaseA genes. **A.** Maximum-likelihood gene phylogeny for *ECP* and *EDN* family members generated using PhyML. Clades representing *ECP* (blue) and *EDN* (green) homologs are indicated. **B.** Diagram showing syntenic regions of the RNaseA gene cluster in humans (top) and white-cheeked gibbons (bottom). Positions of the *ECP* (blue) *EDN* (green) and *RNASE4* (orange) genes are indicated. Rearrangement in the gibbon genome appears to have excised region containing *ECP*, while the *RNASE1* and *RNASE6* genes have undergone translocation to a distal genomic region. **C.** Diagram and crystal structure (PDB: 2LVZ) showing sites subject to positive selection (red) in *ECP*. Sites passed Pr>0.95 cutoff using both Naïve Empirical Bayes and Bayes Empirical Bayes analyses from PAML using both a species phylogeny as well as a maximum-likelihood gene phylogeny. Structure was solved in the presence of a glycosaminoglycan ligand (blue), hypothesized to mediate target cell association. **D.** Summary of predicted gene gain (blue) and loss (red) events of *ECP* in primates.

We next sought to pinpoint amino acid positions in ECP that may have been subject to positive selection in the primate lineage. Given the observed phylogenetic discordance for this gene, we only considered positions that exhibited signatures of positive selection using both a primate species phylogeny as well as a maximum-likelihood *ECP* gene phylogeny (Figure 4C, Supplementary Tables 1, 2, 5, 6-9). Using these criteria we identified numerous sites with strong statistical support for positive selection in ECP (Figure 4C). Rapid evolution of the RNase and glycan-recognition surface of ECP suggests that positive selection may significantly impact ligand affinity or specificity (Figure 4C) (Swaminathan et al. 2001). Many of these rapidly evolving sites also overlap with sites that have been suspected to regulate RNase activity, raising the possibility that this enzymatic function may have been lost, or otherwise modified in divergent primates. Combined with evidence of positive selection acting on *MBP*, our findings suggest that eosinophil granule protein function has been subject to intense selective pressure during primate divergence, perhaps as a consequence of evolutionary conflicts with pathogenic microbes or parasites (Figure 4D).

## DISCUSSION

Our study demonstrates that several core components of the type 2 immune response pathway rapidly evolved in primates, with potential implications for human asthma susceptibility. A previous report also proposes that *IL13* has been subject to positive selection in human populations (Zhou et al. 2004), suggesting that both ancient and ongoing genetic conflicts may be shaping the evolution of this central cytokine. Prior studies of evolutionary arms races intimate that specific molecular interactions between host and microbial factors contribute to signatures of positive selection, as mutations at these interfaces strongly impact comparative fitness (Daugherty and Malik 2012; Duggal and Emerman 2012). Despite its important role in activating Th2-mediated immune responses, we are unaware of any microbe-encoded factors that target IL-13 specifically. Furthermore, we did not detect signatures of selection acting on the IL-13 receptor, IL4R, suggesting that this observed variation in IL-13 is not a product of coevolution with its endogenous receptor (Supplementary Table 10). Numerous pathogens are known to encode interleukin mimics or inhibitors of interleukin signaling (Elde and Malik 2009), suggesting that this is a common and effective strategy of immune evasion. Future studies could assist in identifying specific microbes or factors that specifically target IL-13.

In addition to IL-13, IL-4 is a related cytokine that plays a largely overlapping role in the allergic immune response. While our preliminary survey failed to identify significant signatures of selection in *IL4* among primates, a previous study identified evidence of positive selection acting on multiple sites of *IL4* across diverse mammals (Koyanagi et al. 2010). These findings suggest that IL-4 could be rapidly evolving in other mammalian lineages – indeed, the authors reported functional diversity among rodent IL-4 variants consistent with a recent history of positive selection. Thus, evolutionary pressures acting on particular type 2 immunity factors are likely variable across vertebrate taxa.

Evidence of positive selection acting on multiple eosinophil granule proteins indicates that eosinophil immune function has undergone rapid and recurrent adaptation in primates. These proteins present a “double-edged sword” from the perspective of the host – while they provide potent defensive functions against several types of pathogens and parasites, they also exacerbate tissue damage and inflammation associated with asthma and other autoimmune diseases (Hogan et al. 2008). While these genes provide a transient benefit in the face of selective pressure from pathogens, the fitness cost associated with their toxicity could also drive gene loss events to fixation when pathogen burden is reduced in a population. Conflicts with pathogens or other factors may initially promote gene diversification and step-wise amino acid substitutions for enhanced immune function. Loss of selective pressure from pathogens could subsequently attenuate the fitness gains for this gene, possibly incurring a net cost in the form of autoimmunity. Eosinophil granule genes could therefore provide an informative system to model the dynamics of gene gain, diversification, and loss under changing selective pressures.

In addition to their shared expression within eosinophil granules, MBP and ECP also share a convergent function of cell surface glycan recognition, although the binding spectrum of either protein for glycans from particular microbes or host cells remains unclear. Numerous studies have demonstrated how positive selection can rapidly alter host-pathogen proteinprotein interfaces, although such Red Queen dynamics have not been well-documented for protein-glycan interactions. A recent study found that the typhoid toxin of human-specific *Salmonella* Typhi displays preference for N-acetylneuraminic acid (Neu5Ac), which is abundant on human cells but not those of other mammals, suggesting recent adaptation for human-specific glycan binding. Together these observations suggest that surface glycan recognition could play an broad role in the evolution of host-pathogen interactions within primate hosts as well as other taxa.

A high frequency variant in *ECP* found among humans causes a nonsynonymous change at amino acid position 124 (rs2073342). While this site does not exhibit strong signatures of positive selection across primates, variation at this site has also been linked to the degree of cytotoxicity associated with ECP (Rubin et al. 2009; Rubin and Venge 2013), although its contribution to asthma susceptibility is presently unknown. Future evolution-guided experimental studies of MBP and ECP could assist in determining if and how selection has modulated their distinct protein functions.

The evolutionary causes and significance of allergic immunity continue to be a subject of debate. On the one hand, deleterious allergies and asthma could be interpreted as a simple fitness cost as a trade-off for host immunity. In this view, allergen activation of Th2 immune responses are effectively acting ‘improperly’ against non-harmful stimuli. Alternative hypotheses have proposed that allergies are in fact beneficial and may represent adaptation in response noxious environmental hazards or venoms (Palm et al. 2012). Consistent with this hypothesis, previous work has indicated that allergic responses can mediate clearance of allergens from airway as well as resistance to snake and insect toxins (Tsai et al. 2015). While our findings demonstrate that the type 2 immune response has evolved rapidly and that associated genetic variation may contribute to disease phenotypes, we do not know what pathogens or toxins may have imposed such selective pressure on primate populations. It is notable that certain bacteria native to the upper respiratory tract, such as *Haemophilus influenzae*, are both opportunistic pathogens predicted to drive molecular arms races as well as potential contributors to severe asthma (McCann et al. 2016). While microbial pathogens are often invoked to explain signatures of selection in immunity factors, venom proteins or other biotic toxins could also give rise to such genetic conflicts. Indeed, genes encoding venom proteins have previously been shown to engage in Red Queen dynamics, consistent with this hypothesis (Casewell et al. 2013). Determining the functional implications of genetic variation in type 2 immune response components could provide new inroads for understanding the evolutionary significance of allergic immunity.

It has previously been observed that many human disease-associated substitutions, also termed pathogenic deviations, are fixed or present at high frequency in related mammals (Kondrashov et al. 2002; Barešić et al. 2010). One explanation for these findings is epistasis among compensating mutations that restore “normal” protein activity (Kondrashov et al. 2002; Barešić et al. 2010). Our findings raise the question as to the functional impact of mutations among type 2 immune response genes in other primates. In particular, mutations in *IL13* detected in related primates have known molecular phenotypes associated with enhanced immune signaling and asthma susceptibility in humans (Vladich et al. 2005). While compensating substitutions may be present within these proteins or among interacting factors that alleviate disease phenotypes, it is also possible that these mutations contribute to asthma-like syndromes in non-human animals (Plopper and Hyde 2008; Mueller et al. 2015). Future experimental studies may be able to distinguish between these distinct possibilities.

Our work suggests that natural selection has shaped several protein functions relevant to asthma pathology in primates. This is perhaps best illustrated in the case of *IL13*, where we observe recurrent amino acid substitution of a discrete site across primates linked to asthma susceptibility and increased inflammatory signaling in humans (Wang et al. 2003; Vladich et al. 2005). Understanding patterns of genetic variation among type 2 immunity factors among nonhuman primates could also inform future therapeutic strategies. For example, recurrent loss of *ECP* in primates suggests that there may be a selective advantage to inhibiting this factor pharmacologically. Evolutionary insights could also be critical to improving animal models of autoimmune disease, since genetic differences between humans and other primates could confound, or conversely illuminate, the development of new therapies. Integrating molecular phylogenetic approaches with population genetics and experimentation holds great promise for improving our understanding and treatment of human autoimmune diseases.

## ACKNOWLEDGEMENTS

We are grateful to members of the Elde lab for helpful discussions and comments on the manuscript. We also thank Scott Wong and Adam Geballe for tissue samples and cell lines. This work has been supported by grants from the National Institutes of Health to N.C.E. (R00GM090042) and M.F.B. (F32GM108288; K99GM115822). N.C.E. is the Mario R. Capecchi Endowed Chair in Genetics.

## AUTHOR CONTRIBUTIONS

MFB and NCE conceived the study and wrote the manuscript with assistance from EML. MFB, EML, and HG performed molecular biology experiments. MFB and EML performed phylogenetic analyses with input from NCE.

---

**Supplementary Figure 1.**
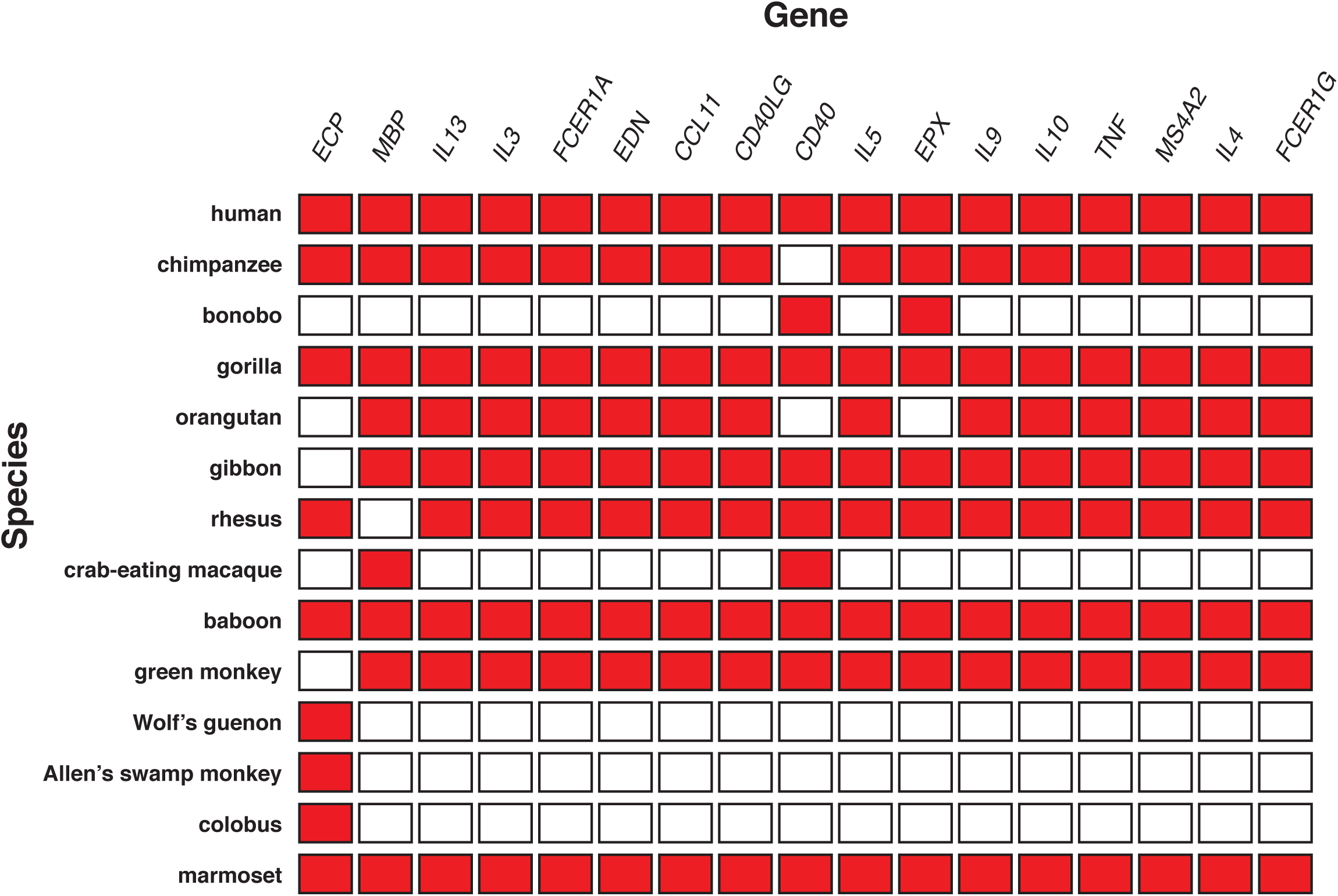
Primate species used in initial screen for evidence of positive selection. Red boxes indicate species for which a gene ortholog was included for phylogenetic analysis.

**Supplementary Figure 2.**
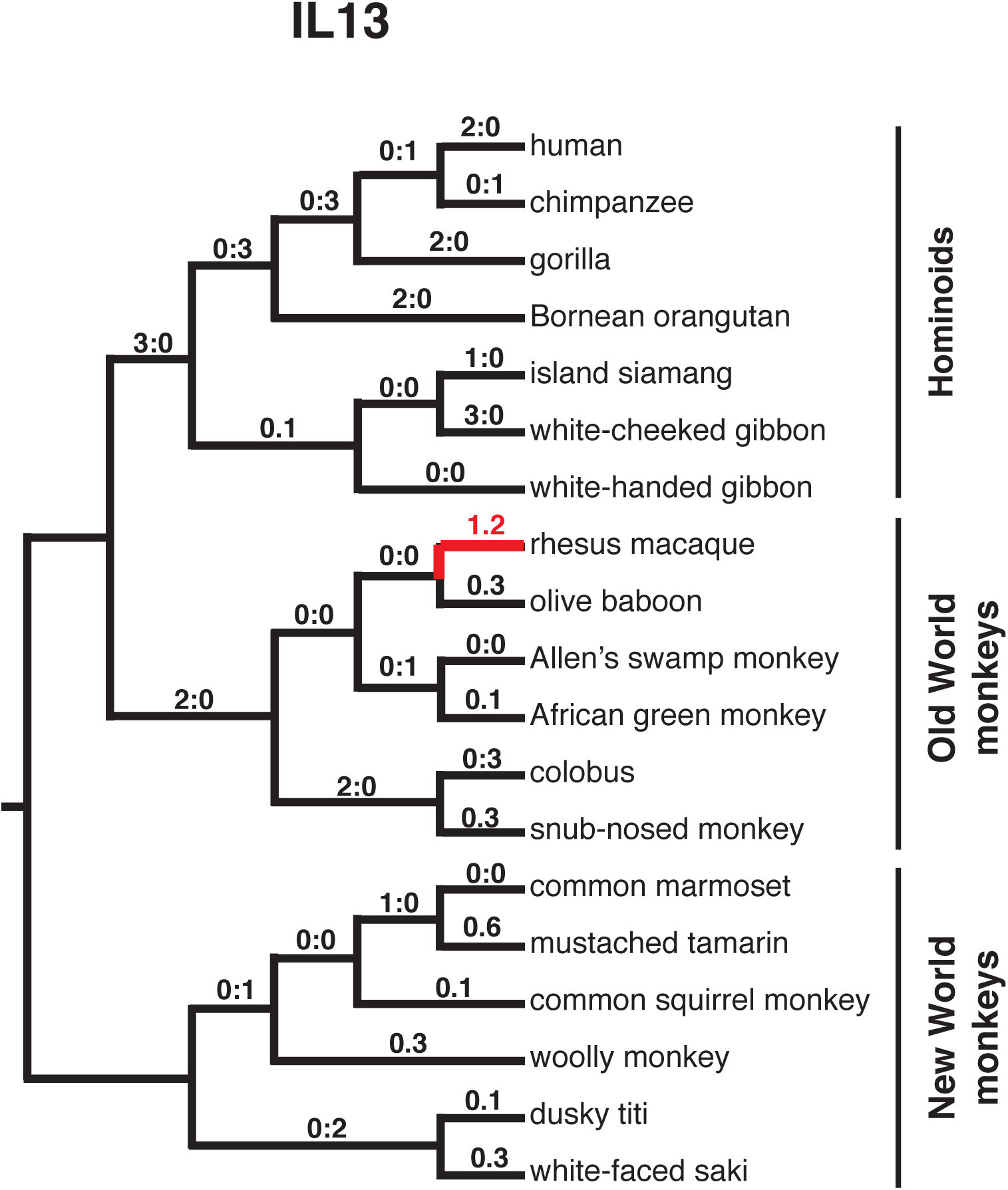
dN/dS ratios for IL13 orthologs in the pirmate lineage as calculated Using PAML. dN/dS ratios greater than 1 are denoted in red.

**Supplementary Figure 3.**
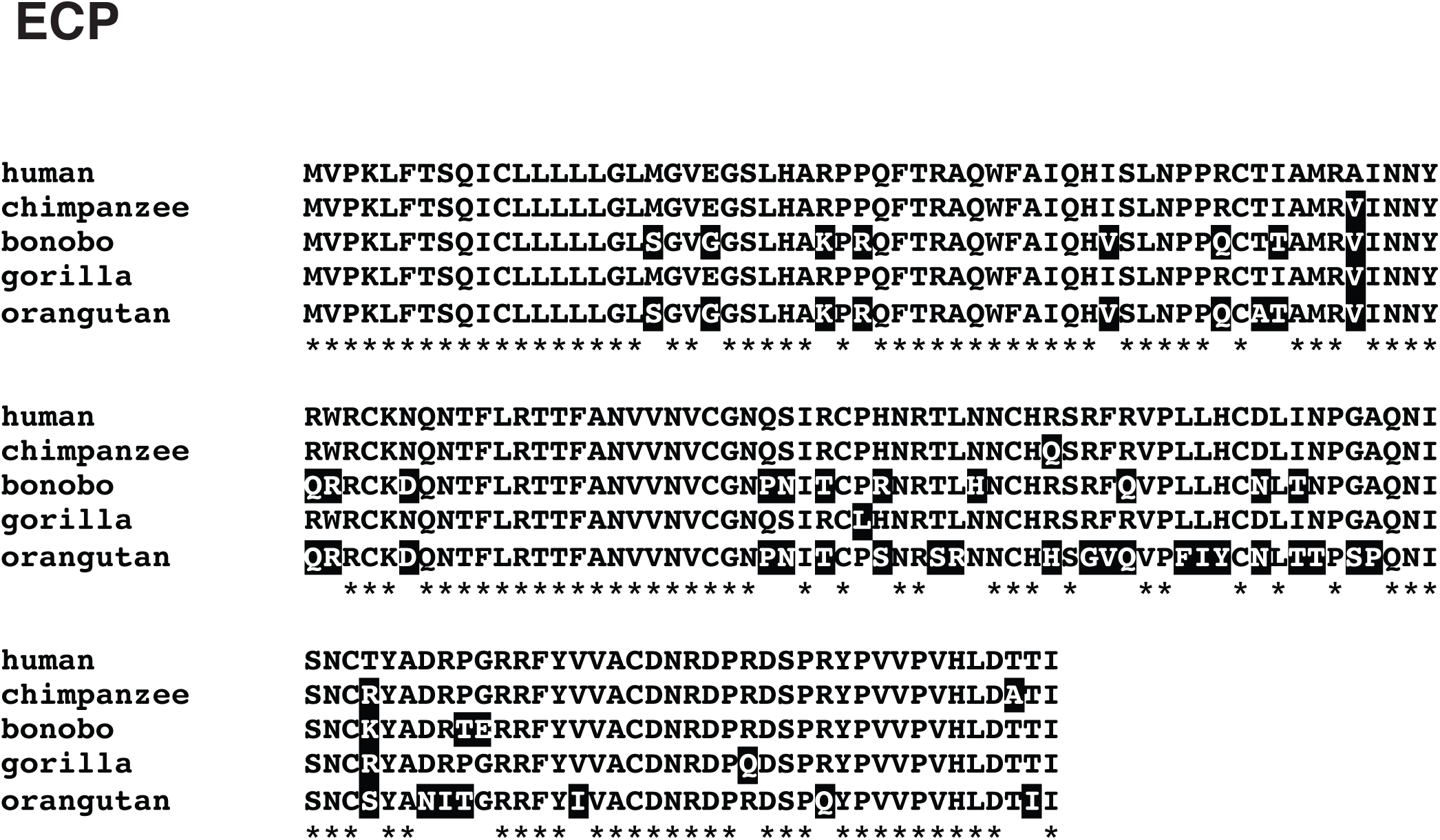
Amino acid alignment of great ape ECP homologs. Divergent sites relative to human ECP are highlighted.

